# Hidden layers of density dependence in consumer feeding rates

**DOI:** 10.1101/2020.08.25.263806

**Authors:** Daniel B. Stouffer, Mark Novak

## Abstract

Functional responses relate a consumer’s feeding rates to variation in its abiotic and biotic environment, providing insight into consumer behavior and fitness, and underpinning population and food-web dynamics. Despite their broad relevance and long-standing history, we show here that the types of density dependence found in classic resource- and consumer-dependent functional-response models equate to strong and often untenable assumptions about the independence of processes underlying feeding rates. We first demonstrate mathematically how to quantify non-independence between feeding and consumer interference and between feeding on multiple resources. We then analyze two large collections of functional-response datasets to show that non-independence is pervasive and borne out in previously-hidden forms of density dependence. Our results provide a new lens through which to view variation in consumer feeding rates and disentangle the biological underpinnings of species interactions in multi-species contexts.

## Introduction

Functional responses are a critical component in our understanding of consumer-resource interactions. Since the pioneering work of Holling (1959a;b; 1965), numerous researchers have used manipulative and observational experiments to elucidate the empirical ways in which varied biotic and abiotic factors alter consumer feeding rates in diverse biological contexts (e.g., DeLong, 2014; Preston *et al*., 2018; Uiterwaal & DeLong, 2020). In parallel, researchers have proposed a large suite of models to characterize functional responses mathematically (Abrams & Ginzburg, 2000; Jeschke *et al*., 2002; Gentleman *et al*., 2003; Koen-Alonso, 2007; Arditi & Ginzburg, 2012), with emphasis increasingly being placed on the evaluation of their statistical performance and empirical relevance (Skalski & Gilliam, 2001; Jeschke *et al*., 2004; Novak *et al*., 2017; Uiterwaal & DeLong, 2020).

Though they differ in their finer details, one assumption that is common to virtually all functional-response models is that feeding rates will exhibit density-dependent variation. The agents of this density dependence provide a convenient classification scheme: on one hand, we have “resource-dependent” or “consumer-dependent” models (whereby a focal consumer’s feeding rate is either determined by resource abundance alone, or by the abundances of resources and consumers together; Abrams & Ginzburg, 2000); on the other, we have “single-resource” versus “multi-resource” models (whereby focal consumers are assumed to be specialists feeding on a single resource, or generalists whose feeding rates on non-focal resources could influence their feeding rate on the focal resource; Murdoch, 1973). For example, the wellknown Holling Type II functional-response model, *F* (*N*) = *aN/*(1+*ahN*), is a single-resource dependent model since the per capita feeding rate *F* saturates only as a function of increasing resource density *N* (Holling, 1959b). In this model, the rate of saturation is determined by the “attack rate” *a* and the “handling time” *h*, the latter of which imposes an ever greater cost to the consumer as the rate with which it encounters (and captures) resources increases with resource density.

Despite the ubiquity of density dependence in functional-response models, the parameters that control how species densities affect variation in feeding rates are rarely themselves considered to be explicit functions of species’ densities (Abrams, 1982; Kéfi *et al*., 2012). Although the Holling Type III model (Holling, 1959b) may be interpreted as assuming that the attack rate is a linearly increasing function of resource density *N*, it is far less common to allow handling time to also depend on *N* (but see Abrams, 1990; Okuyama, 2010). And yet, while almost all resource-dependent and consumer-dependent models consider feeding rate saturation to be a function of (i) increasing resource density via handling times and (ii) increasing consumer density via conspecific interference, neither handling times nor interference rates are themselves considered to depend on consumer or resource densities, respectively.

Broadly-speaking, density dependence of interaction-rate parameters leads to interaction modifications (Wootton, 1993; Adler & Morris, 1994; Goudard & Loreau, 2008) reflecting indirect effects (Wootton, 1994; Okuyama & Bolker, 2007; Abrams & Cortez, 2015), trait- and behavior-mediated effects (Beckerman *et al*., 1997; Peacor & Werner, 2001; Werner & Peacor, 2003; Toscano & Griffen, 2014), and other forms of non-additivity or higher-order effects (Mayfield & Stouffer, 2017; Letten & Stouffer, 2019; Kleinhesselink *et al*., 2019). Though these phenomena have long been recognized as being biologically widespread (Abrams, 1983; Strauss, 1991; Levine *et al*., 2017), there are multiple explanations for why they remain underrepresented in the functional-response literature and why their potential importance for consumer feeding rates has not been empirically addressed. Among these reasons are the high logistical costs associated with even the simplest of functional-response experiments, with statistical insight into additionally-assumed parameters requiring ever more treatment levels and greater amounts of replication (Beck & Arnold, 1977; Bolker, 2008). Researchers are also well-justified in wishing to avoid unnecessary increases in model complexity that complicate mathematical analyses and can lead to over-fit statistical models (Rissanen, 1996; Myung *et al*., 2000; Burnham & Anderson, 2002). A more fundamental challenge, however, is that it is far easier to add potentially unnecessary new terms to a model than it is to provide a biological rationale for why they should be included (Abrams, 1997; Ginzburg & Jensen, 2004; Otto & Day, 2007; Guimerà *et al*., 2020). This represents a general problem for the functional-response literature because it lacks a general perspective from which to biologically motivate such terms.

To address this challenge, we provide a mathematical analysis to demonstrate how these under-studied density-dependent terms can emerge from classic consumer functional-response models. We focus our analysis on two broadly-studied scenarios: First, we consider the case of multiple conspecific consumer individuals foraging on a single resource species. In this context, we derive new models that generate a spectrum of emergent consumer-interference effects that have not been previously described. Second, we consider the case of a single consumer individual foraging on two different resource species. In this context, we derive new models that generate a spectrum of emergent effects between resource species that have also not been previously recognized. To assess the empirical relevance of these new functional-response models and thereby motivate targeted experimental designs in the future, we then fit them to two large collections of published functional-response data representing consumer identities that range from wolves to ciliates. Our analysis provides evidence for the widespread prevalence of unrecognized density-dependent effects in many existing functional-response experiments.

## Mathematical analysis

We first show how links between the various processes that underlie feeding rates can create novel functional forms for feeding rate variation; specifically when the rates of consumer interference and/or consumption of different resources are, or are not, independent of each other. Our analysis demonstrates why non-independence leads directly to new functional-response models containing “density dependence of interaction-rate parameters” (e.g., handling times that are explicit functions of consumer densities). For simplicity and to better relate to the prevailing literature, we will generally refer to consumers as predators and resources as prey. However, we will subsequently use our data analysis to show that the scenarios described apply to consumer–resource interactions more broadly.

### Single-resource consumer dependence

We first consider how interactions between conspecific predators act to change their own per capita feeding rate. One of the simplest models that includes such interactions by allowing for both resource and consumer density dependence is the single-resource consumer-dependent Beddington–DeAngelis functional response (Beddington, 1975; DeAngelis *et al*., 1975). This model takes the form

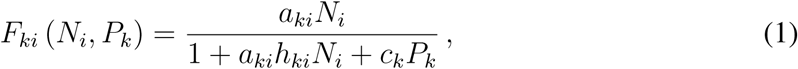

where *F*_*ki*_ is the feeding rate of predators *k* consuming prey *i* (with dimensions of [(prey consumed per predator) per time available]), *N*_*i*_ is the number of prey *i* available, *P*_*k*_ is the number of predators *k, a*_*ki*_ is the attack rate (with dimensions of [(prey consumed per prey available) per time available]), *h*_*ki*_ is the handling time (with dimensions [time handling per prey consumed]), and *c*_*k*_ is the strength of interference between predators (with dimensions [(time interfering per time available) per predator interfering]). When *P*_*k*_ represents a count rather than a density of predators, as is common in experimental settings, *P*_*k*_ is replaced by (*P*_*k*_ − 1) because a predator individual cannot interfere with itself. Note that the 1 in the denominator is dimensionless for the same reason that time cancels out in the dimensions of interference strength *c*_*k*_. We have also refrained from including dimensions of area or volume in *a*_*ki*_ or *c*_*k*_ because they have no impact on our subsequent data analysis (but see Uiterwaal & DeLong, 2020).

A related model, the Crowley–Martin functional response (Crowley & Martin, 1989), takes the form

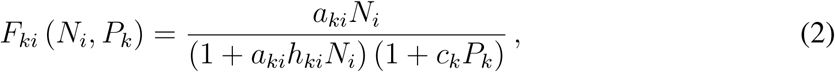

where all parameters are defined precisely as above. Whereas the Beddington–DeAngelis model is interpreted as characterizing predators that only interfere when searching for prey, the Crowley–Martin model is interpreted as characterizing predators that interfere both when searching for and when handling prey.

Focusing on the denominators of Eqs. (1–2), the key mathematical difference between the Beddington–DeAngelis and Crowley–Martin models is an additional term in the latter that varies with the product of *N*_*i*_ and *P*_*k*_. We could therefore instead rewrite both models as

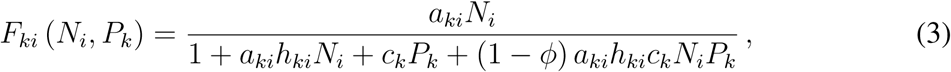

where *ϕ* is a dimensionless parameter that controls the strength of this *N*_*i*_*P*_*k*_ term. Written in this way, we immediately recover the Beddington–DeAngelis model when *ϕ* = 1 and the Crowley–Martin model when *ϕ* = 0.

### Understanding the parameter *ϕ*

We can conceptualize the role of *ϕ* in creating density-dependent functional-response parameters in various ways. For example, we could rearrange the denominator of Eq. (3) to instead give

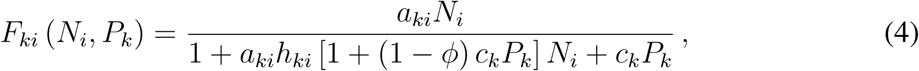

in which case the implied interpretation is that *ϕ* influences the extent to which handling time *h*_*ki*_ is a function of the abundance of interfering predators *P*_*k*_. That is, *h*_*ki*_ is only independent of interfering predators when *ϕ* = 1. One could equivalently rearrange Eq. (3) to give

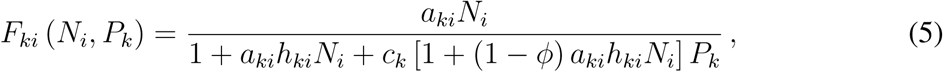

in which case the implied interpretation is that *ϕ* influences the extent to which interference strength *c*_*k*_ is a function of the abundance of available prey *N*_*i*_. That is, *c*_*k*_ is only independent of available prey when *ϕ* = 1. From a phenomenological perspective, it is important to recognize that both viewpoints are equally correct—as are combinations of them—as they are mathematically identical. Stepping back a bit, this implies that the most parsimonious role of *ϕ* is as an indicator that neither prey dependence—as captured with handling time *h*_*ki*_—nor predator dependence—as captured with interference strength *c*_*k*_—can be properly measured independent of the other.

To clarify the origin of this dependence of *h*_*ki*_ and *c*_*k*_ on species densities, we extend a derivation previously presented by Crowley & Martin (1989). Rather than describe predator– prey functional responses phenomenologically, as formulated above, those authors described how the observed feeding rate *F*_*ki*_ relates to an implicit, unmeasured interference rate *I*_*k*_:

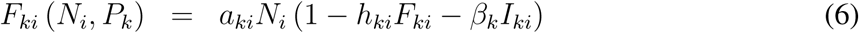

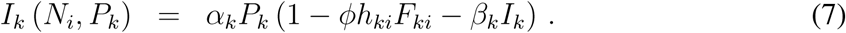

Upon algebraically solving Eqs. (6–7) for *F*_*ki*_, we immediately obtain Eq. (3) with interference strength *c*_*k*_ = *α*_*k*_*β*_*k*_. The two new parameters are an “interference rate” *α*_*k*_ (akin to the attack rate and with dimensions [(predator interfered per predator interfering) per time available]), and an “interference time” *β*_*k*_ (akin to handling time and with dimensions [time interfering per predator interfered]). The parenthetical term of each equation therefore corresponds to the proportion of total time available for attacking and the proportion of total time available for interfering, respectively. That is, consistent with the definitions of *h*_*ki*_ and *β*_*k*_, time for searching in Eq. (6) is reduced by time spent handling and time spent interfering. Similarly, time available for interfering in Eq. (7) is also reduced by interfering. In contrast, whether and how time spent handling influences the realized rate of interference is explicitly determined by the value of *ϕ*.

Expressed in this way, we can shift from a generic parameter *ϕ* and formally define 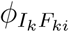 as the parameter capturing how the predator’s feeding rate *F*_*ki*_ alters its realized conspecific interference rate *I*_*k*_. As described verbally above, predators can interfere while searching but cannot interfere while feeding in the Beddington–DeAngelis model (Eq.1), implying 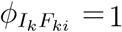. In the Crowley–Martin model (Eq.2), predators interfere both while searching and while feeding, implying 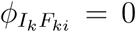. Crowley & Martin (1989) referred to these as distraction and distraction-free models, respectively. Beyond these two cases, note that *any* value of 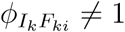 leads to the appearance of a “higher-order” term in the denominator that involves the product of both prey and predator densities, *N*_*i*_*P*_*k*_. Parameter 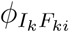 causes feeding rates to *decrease* with increasing *N*_*i*_*P*_*k*_ whenever 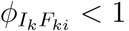, and causes feeding rates to *increase* with increasing *N*_*i*_*P*_*k*_ whenever 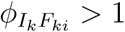 (Fig. 1a).

**Figure 1:**
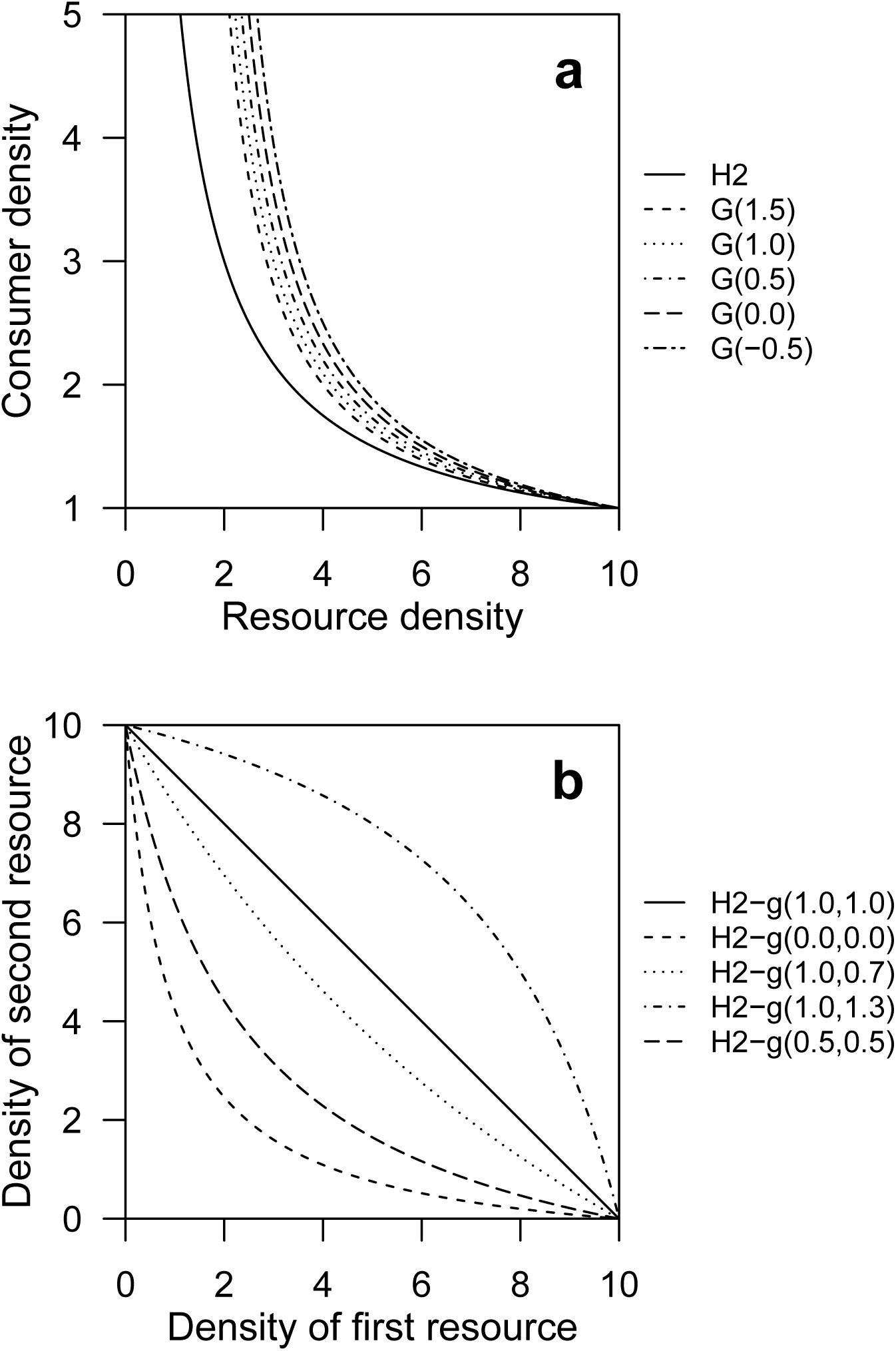
**a**, The effect of the parameter 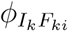 on feeding rate *F*_*ki*_. We show isoclines of constant consumption for the Holling Type II model (H2) and the generalized resource- and consumer-dependent model introduced here (Eq. 3). Each line corresponds to *P*_*k*_*F*_*ki*_ ≡ 1 with the values of 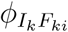 specified in the legend, and fixed values of attack rate *a*_*ki*_ = 0.2, handling time *h*_*ki*_ = 0.5, and interference *c*_*k*_ = 0.25. As 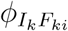 shifts from 1.5 (short-dashed line) to −0.5 (dot-dashed line), more and more consumers are required to achieve equivalent consumption. Note that G(1) and G(0) correspond to the Beddington–DeAngelis and Crowley–Martin models, respectively. **b**, The effect of the parameters 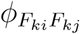 and 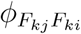 on total feeding rate *F*_*k*_ = *F*_*ki*_ + *F*_*kj*_. We show isoclines of constant consumption for the generalized multi-resource dependent model introduced here (Eq. 10). Each line corresponds to *P*_*k*_*F*_*k*_ ≡ 1 with the values of 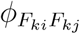 and 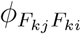 specified in the legend and *P*_*k*_ = 1. Both resources are otherwise equivalent (i.e., *a*_*ki*_ = *a*_*kj*_ = 0.5 and *h*_*ki*_ = *h*_*kj*_ = 0.5). When 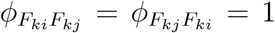 (solid line), resources are perfectly substitutable; when 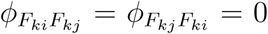, consumers require fewer resources to achieve equivalent feeding rates (short-dashed line); and when 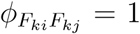 and 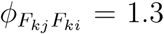, consumers require more resources to achieve equivalent feeding rates (dot-dashed line). Note that H2-g(1, 1) corresponds to the multi-resource Holling Type II functional response (H2-m).

### Multi-resource dependence

We follow a similar methodology for multi-resource dependence—as occurs when a single predator is feeding on two different prey species—to measure the effect that feeding on one prey species has on the predator’s rate of feeding on the second prey species (and vice versa). Similar to Eqs. (6–7), we define the feeding rates on prey *i* and prey *j* as

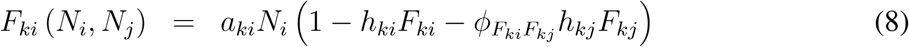

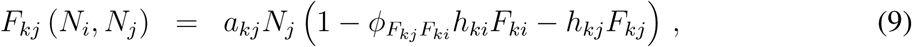

where 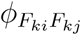 represents the extent to which feeding on *j* impacts feeding on 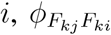 represents the extent to which feeding on *i* impacts feeding on *j*, and all other parameters are interaction-specific versions of the quantities already defined above (e.g., *a*_*ki*_ and *a*_*kj*_ vs. only *a*_*ki*_). Hence the parenthetical term of each equation corresponds to the proportion of total time available for attacking *i* (respectively *j*) after accounting for time spent handling both prey.

With algebraic manipulation of Eqs. (8–9), we can solve for each of the two feeding rates to give

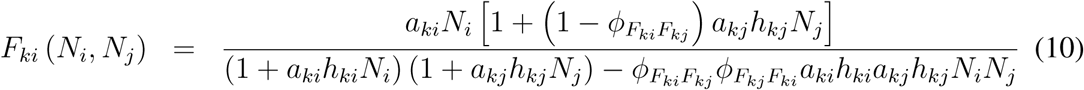

for the predator’s feeding rate on *i*. (An equivalent expression for the predator’s feeding rate on prey *j* can be obtained by swapping all *i*’s for *j*’s and vice versa.)

As was the case for predator interference, *any* values of 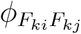 and 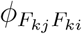 such that their product 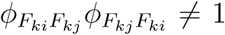 lead to the appearance of the additional “higher-order” term in the denominator of Eq. (10) involving the product of both prey densities, *N*_*i*_*N*_*j*_. For the multi-resource dependence case, the density of the second prey also appears in the numerator of Eq. (10). As a direct consequence, non-independence between feeding on both prey species as captured by the parameters 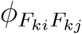 and 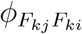 can lead to density-dependent changes in feeding of the sort expected when predators exhibit prey-switching behavior (Supplementary Material).

In order to better elucidate the behavior of these expressions for feeding rates, it is useful to explore three limiting cases and the resulting forms for *F*_*ki*_. First, consider a scenario in which 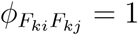 and 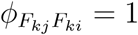, implying that predators cannot search for any prey while handling either, as is likely true for intertidal whelks that feed on sessile prey (e.g., Novak *et al*., 2017). Here the dependence on the “other” prey vanishes from the numerators of both feeding-rate expressions and the higher-order term cancels within both denominators. As a result, we are left with the standard “multi-resource Holling Type II functional response” (Murdoch, 1973; Koen-Alonso, 2007) given by

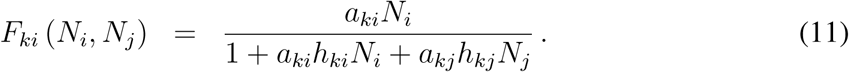

Second, consider the scenario where 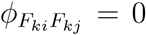, implying that the feeding rate on *j* has no impact on the feeding rate on *i* as might occur if prey are only available during different times of day (e.g., Forrester *et al*., 1994). In this case, the higher-order term vanishes from the denominator in *F*_*ki*_ irrespective of the value of 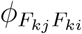. Moreover, the potential dependence on *N*_*j*_ can be factored out since identical expressions of the form 1 + *a*_*kj*_*h*_*kj*_*N*_*j*_ appear in both the numerator and denominator. Reassuringly, we are left with a single-resource Holling Type II functional response for *F*_*ki*_ that is independent of the abundance of prey *j* (Holling, 1959a;b); namely we obtain

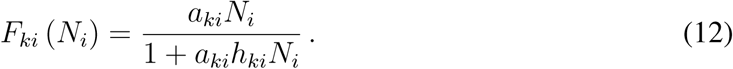

Third, consider what happens when 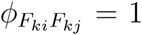 and 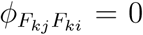, implying that a predator handling *j* cannot attack *i* but a predator handling *i* could still attack *j*. This scenario could arise when prey differ dramatically in size (e.g., Kalinkat *et al*., 2011). Under these conditions, feeding on *i* behaves as

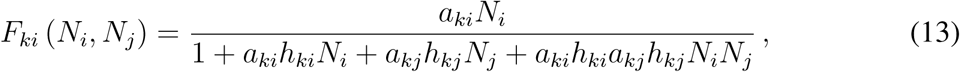

which shows systematic variation for any abundance of *j* and includes the higher-order term in the denominator. Compared to first two limiting cases, feeding *F*_*ki*_ is lowest in this third scenario (for equivalent values of the attack rates and handling times). This is because increased feeding *F*_*kj*_ on *j* acts to decrease the time available for feeding on *i*. Delineating values of 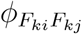 and 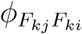 that always lead to increased or decreased feeding rates is more complicated than for single-resource consumer dependence since both rates are a combination of both parameters. Moreover, two given values of 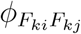 and 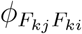 could hypothetically lead to a decrease in *F*_*ki*_ while still increasing the *total* feeding rate *F*_*k*_ = *F*_*ki*_ + *F*_*kj*_ (Fig. 1b).

## Methods

### Data compilation

Our mathematical analysis indicates that *any* non-independence between processes such as feeding and interference can induce higher-order, non-additive terms in the denominator of common functional-response models. We therefore next aimed to determine whether there is empirical support for such non-independence and the inclusion of the parameters 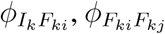, and 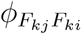 when modeling observed feeding rates. Focusing on the two specific scenarios discussed above, we searched the literature to obtain two different types of empirical datasets. First, single-resource consumer-dependence datasets designed specifically to assess consumer interference; these consisted of feeding rates as a function of variation in prey and predator abundances, or parasitism rates as a function of variation in host and parasitoid abundances (Table S1). Second, multi-resource dependence datasets designed to assess the dependence of consumer feeding rates on the availability of two alternative resources; these universally consisted of feeding rates for single predator individuals as a function of variation in the abundances of two prey (Table S2). When possible, we obtained the original data from the authors. Otherwise, we extracted (i) data points or (ii) means and their associated uncertainties from the publication in tables by hand or figures using GraphClick (2010).

### Single-resource consumer-dependent models

We considered five different functional-response models when examining the consumer-interference datasets (Table 1): the resource-dependent, consumer-independent Holling Type I and Holling Type II models (Holling, 1959a;b), the resource- and consumer-dependent Beddington–DeAngelis (Beddington, 1975; DeAngelis *et al*., 1975) and Crowley–Martin (Crowley & Martin, 1989) models, and our new resource- and consumer-dependent model with the additional parameter 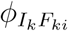 (Eq. 3).

**Table 1:**
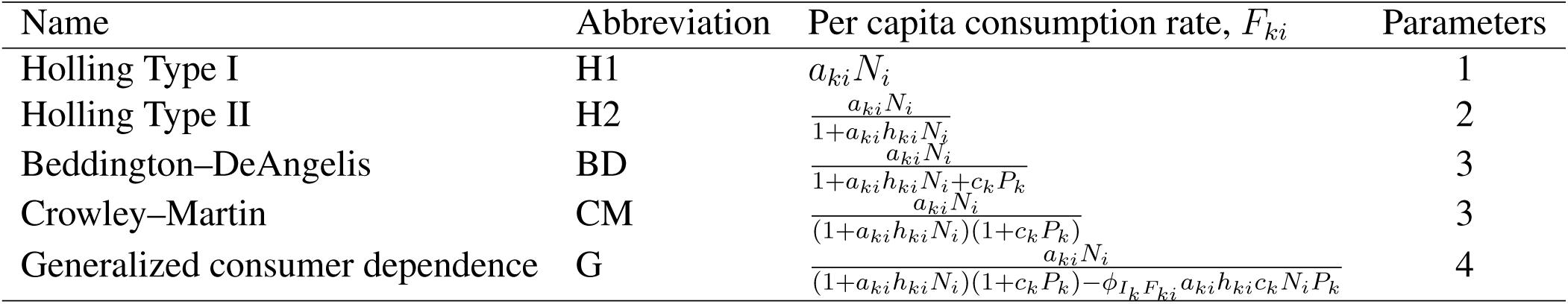
The five models we considered describing single-resource consumer-dependent consumption. These expressions for consumer per capita consumption rates correspond to the instantaneous consumption rate for both replacement and non-replacement studies.

### Multi-resource dependent models

We considered four different functional-response models when examining the multiple-resource datasets (Table 2): the Holling Type I functional responses that arise when *h*_*ki*_ = *h*_*kj*_ = 0, the Holling Type II functional responses that arise when 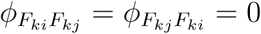 (Holling, 1959a;b), the multi-resource Type II functional responses that arise when 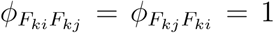 (Murdoch, 1973), and our generalized multi-resource Type II functional responses that emerge when both 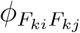 and 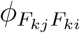 are free parameters (Eq. 10).

**Table 2:**
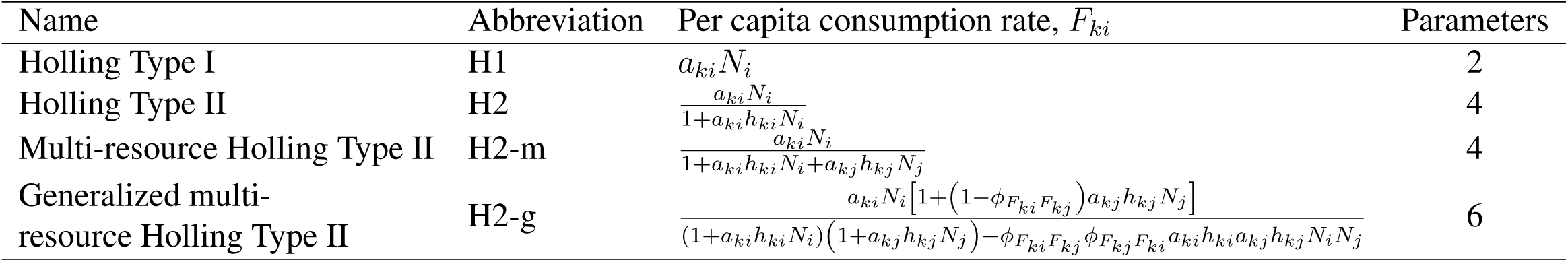
The four models we considered describing multi-resource dependent consumption. These expressions for consumer per capita consumption rates correspond to the instantaneous consumption rate for both replacement and non-replacement studies. As there are two resources under consideration, the number of parameters corresponds to the total number across *both* resources.

### Biological and statistical constraints to the parameters

There is a particularly important detail to consider when fitting Eqs. (3) & (10) to data. In the preceding mathematical descriptions, we primarily focused on examples in which the various parameters 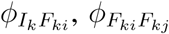, and 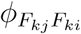 took values of 0 or 1. We did so to build on the intuition behind the distraction and distraction-free interpretations of predator behavior (Crowley & Martin, 1989), and to relate our generalized models to pre-existing functional-response models. This notwithstanding, the values these parameters may take on are not restricted to this 0 to 1 region. Instead, their values depend on the extent to which they generate biologically plausible (or implausible) feeding rate behavior. This may be understood as follows.

When considering the processes involved, the most fundamental constraints are that the rate of predator *k* feeding on prey *i* (*F*_*ki*_), the rate of predator *k* feeding on prey *j* (*F*_*kj*_), and interference rate of predator *k* (*I*_*k*_) in Eqs. (6–9) must each remain greater than or equal to zero. This means, for example, that the statistical best-fit value of 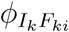 for a given single-resource consumer-dependence dataset could be 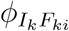 < 0 if *β*_*k*_ were sufficiently large, or could be 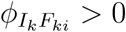 if *h*_*ki*_ were sufficiently small. This contrasts to parameters such as handling time that are directly constrained by their explicit interpretation (e.g., the time associated with handling a prey cannot be negative). Similar arguments hold for 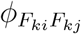 and 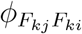 in the context of multiple resources. While an absence of constraints on 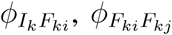, and 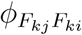 does not impact the mathematical derivation performed above, it does influence their statistical inference as outlined below. It also impacts a model’s ability to generate biologically plausible “out-of-sample” predictions. That is, a large positive value of 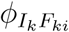 may be consistent with a given dataset while also predicting implausible, negative interference rates for larger-than-observed consumer or resource abundances (see also Novak & Stouffer, 2020).

### Parameter inference

Given each empirical dataset, we determined the best-fit parameter values for each functional-response model using a two-step procedure. In the first step, we used the global-optimization algorithm *sbplx* from the *nloptr* package (Johnson, 2020) in R (R Core Team, 2020) to obtain an estimate of the maximum-likelihood parameter values. In the second step, we passed the optimal parameter values identified by *sbplx* to the *mle2* function from the *bbmle* package (Bolker & R Development Core Team, 2020) to search for local improvements and assess model convergence. The likelihood being optimized was determined by the dataset’s experimental design (Supplementary Material), and all handling times, attack rates, and interference rates were constrained to be positive. We allowed the values of 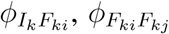, and 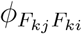 to vary freely as long as predicted mean feeding rates remained greater than or equal to zero.

When a fit converged (i.e., when the maximum-likelihood parameter values were identified), we first attempted to estimate parameter uncertainty via their 68% profile confidence intervals—which roughly correspond to standard error—using the *confint* function from *bbmle*. On occasion, this approach failed because, although the optimization converged, the likelihood surface was nearly flat around the optimum. This is usually indicative of the model being over-parameterized (Gill & King, 2004), the occurrence of which was not altogether surprising in our context since none of the experiments were designed for the purpose of fitting our most complex functional-response models. In these instances, we used the quadratic approximation of the parameter standard errors provided directly by *mle2* as our estimate of parameter uncertainty.

When we could only obtain a dataset as means and associated uncertainties, we simulated 250 parametrically bootstrapped datasets with a sample size equivalent to that of the original dataset and then inferred the best-fit parameter values to each of these (Supplementary Material). We then performed the same two-step parameter fitting process separately for each of these simulated datasets. We treated each parameter’s median value across these 250 separate fits as its point estimate. As an estimate of its uncertainty, we used the central 68% interval of the 250 values as this corresponds to *±*1 standard deviation for a normal distribution.

### Model assessment and model comparison

After fitting the parameters of the various functional-response models, we focused on two primary ways in which the data could lend support to the processes captured by the parameters 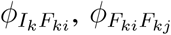, and/or 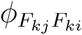. The first came from comparing *AIC* (Akaike Information Criterion) across the various candidate models. Smaller values of *AIC* provide an indication of a better out-of-sample prediction error conditional on model complexity. The second was provided by our aforementioned ability to infer maximum-likelihood values for these parameters that differed from values of 0 or 1 with well-defined, and relatively small, estimates of uncertainty. Even when a model is not the most parsimonious among a set of alternative models, well-defined estimates of parameter uncertainty are still a good indication that it provides a robust description of the data-generating process (Beck & Arnold, 1977; Reichert & Omlin, 1997; Gill & King, 2004).

## Results

### Consumer interference

We obtained 77 single-resource consumer-dependence datasets with which to infer the effect of feeding on consumer interference (Table S1). This included 61 datasets with predator consumers and 17 datasets with parasitoid consumers. In total, we obtained 44 datasets in the form of raw data and 33 datasets in the form of means and associated uncertainties. On average, the datasets consisted of 120 replicate feeding observations (min: 10, max: 528, median: 80).

As judged by *AIC*, our generalized consumer-dependent model including 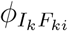 (Eq. 3) was ranked first for 42 (55%) datasets and tied for first (i.e., was within 2 *AIC* units) for an additional 24 (31%) datasets (Fig. 2). We obtained qualitatively similar results using the alternative information criteria *AIC*_*c*_ and *BIC* (Fig. S1). Maximum-likelihood point estimates of 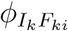 (i.e., the effect of feeding on interference) varied considerably across the datasets (Fig. 2). Well over half (47 out of 77) of the datasets provided point estimates that were less than or equal to one, implying that handling times and/or consumer interference *increased* as the product of resource and consumer abundances increased. The uncertainties of these 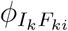 point estimates overlapped only the Beddington–DeAngelis model 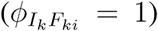 for 15 datasets, overlapped only the Crowley–Martin model 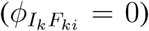 for 11 datasets, and overlapped both models for 23 datasets. This overlap was largely consistent with instances where the generalized model was judged equivalent to simpler models based on *AIC*. A full 28 datasets had uncertainties that were not consistent with *any* pre-existing model: 4 datasets fell entirely in the region 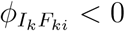, 10 datasets fell exclusively between the two models (e.g., consistent with “partial” distraction of consumers), and 14 datasets fell entirely in the region 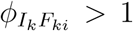. Overall, the uncertainties of only 34 datasets (43%) were consistent with the idea that interference and feeding were independent of each other (i.e., 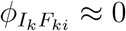).

**Figure 2:**
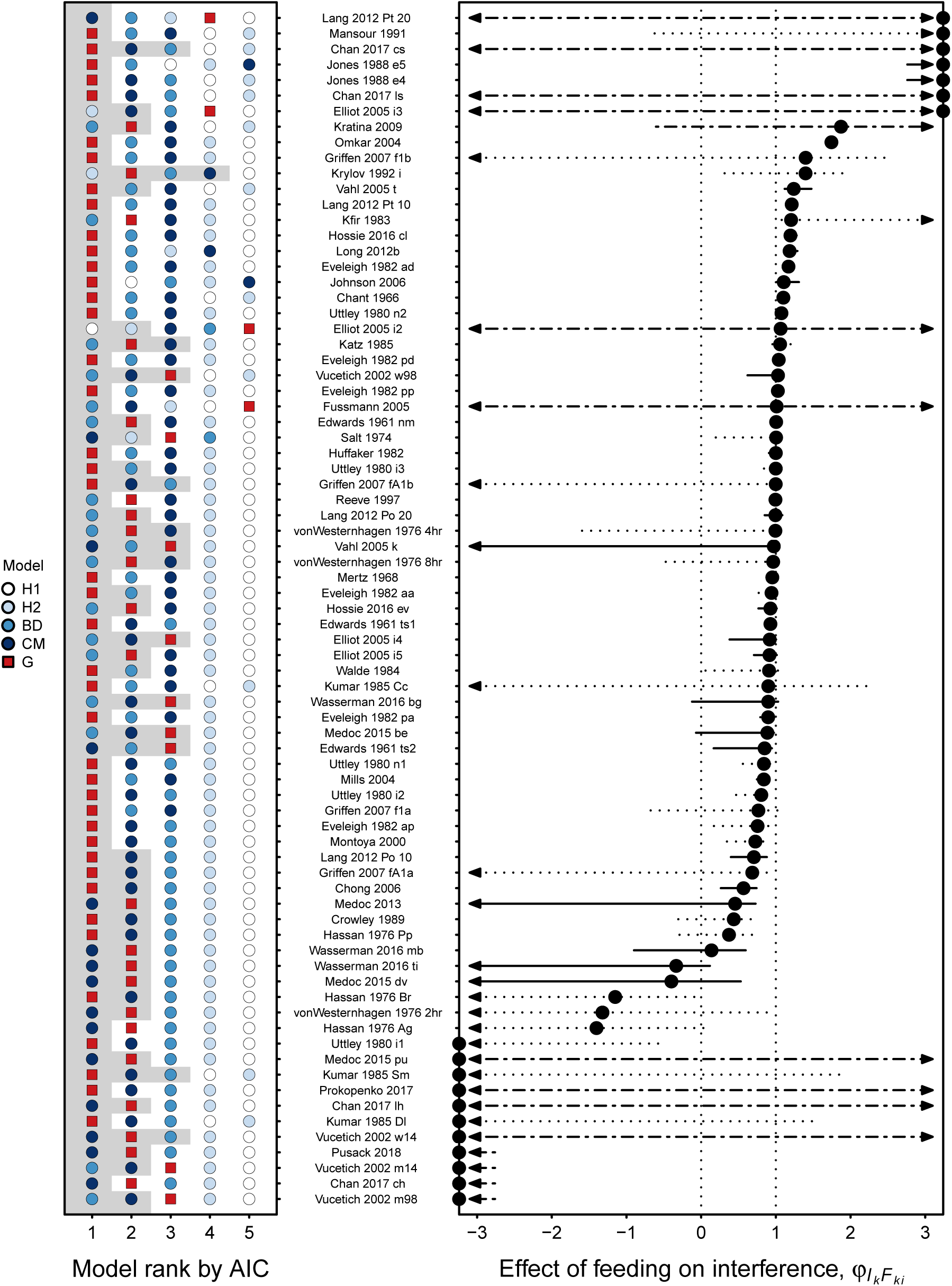
The effect of feeding on consumer interference as estimated across 77 single-resource consumer-dependence datasets. One the left, we show the rank-order performance of functional-response models as judged by *AIC*, with rank 1 indicating the best model and rank 5 indicating the worst. The gray region indicates models with statistically equivalent support (i.e., Δ*AIC* < 2). The red square is the generalized consumer-dependent model introduced here (G). (See Table 1 for all model abbreviations.) On the right, we show the estimated mean and uncertainty for the effect of feeding on interference (i.e., 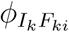). The vertical line at 0 corresponds to the Crowley–Martin model (CM), and the vertical line at 1 corresponds to the Beddington– DeAngelis model (BD). Point estimates outside the plot region are indicated with circles on the plot border. Line types indicate the method for estimating uncertainty: solid for profile confidence intervals, dot-dashed for quadratic approximation, and dotted for bootstrapped data. Uncertainty estimates that are fully or partially beyond the plot region are indicated by arrows. The large uncertainty for some datasets is most likely due to poor parameter identifiability.

### Multiple resources

We obtained 30 multi-resource dependence datasets with which to infer the effect that feeding on one prey has on feeding on another and vice versa (Table S2). The consumers of all of these were predators. This included 15 datasets in the form of raw data and 15 datasets in the form of means and associated uncertainties. On average, the datasets consisted of 135 replicate pairs of feeding observations (min: 37, max: 290, median: 116).

As judged by *AIC*, our generalized multi-resource Holling Type II model including both 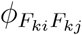 and 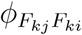 (Eq.10) was ranked first for 20 (67%) multiple-resource datasets and tied for first for an additional 1 (3%) dataset (Fig. 3). We obtained qualitatively similar results using the alternative information criteria *AIC*_*c*_ and *BIC* (Fig. S2). Maximum-likelihood point estimates of 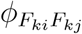 and 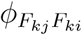 varied considerably across the different datasets (Fig. 3), including estimates indicative of (i) feeding rates being unaffected by the non-focal resource (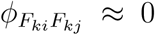 and 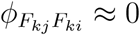), (ii) feeding on one resource completely precluding feeding on the other (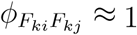 and/or 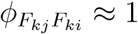), (iii) feeding on one resource only partially precluding feeding on the other (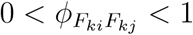 and 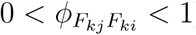), and (iv) almost all combinations of these.

**Figure 3:**
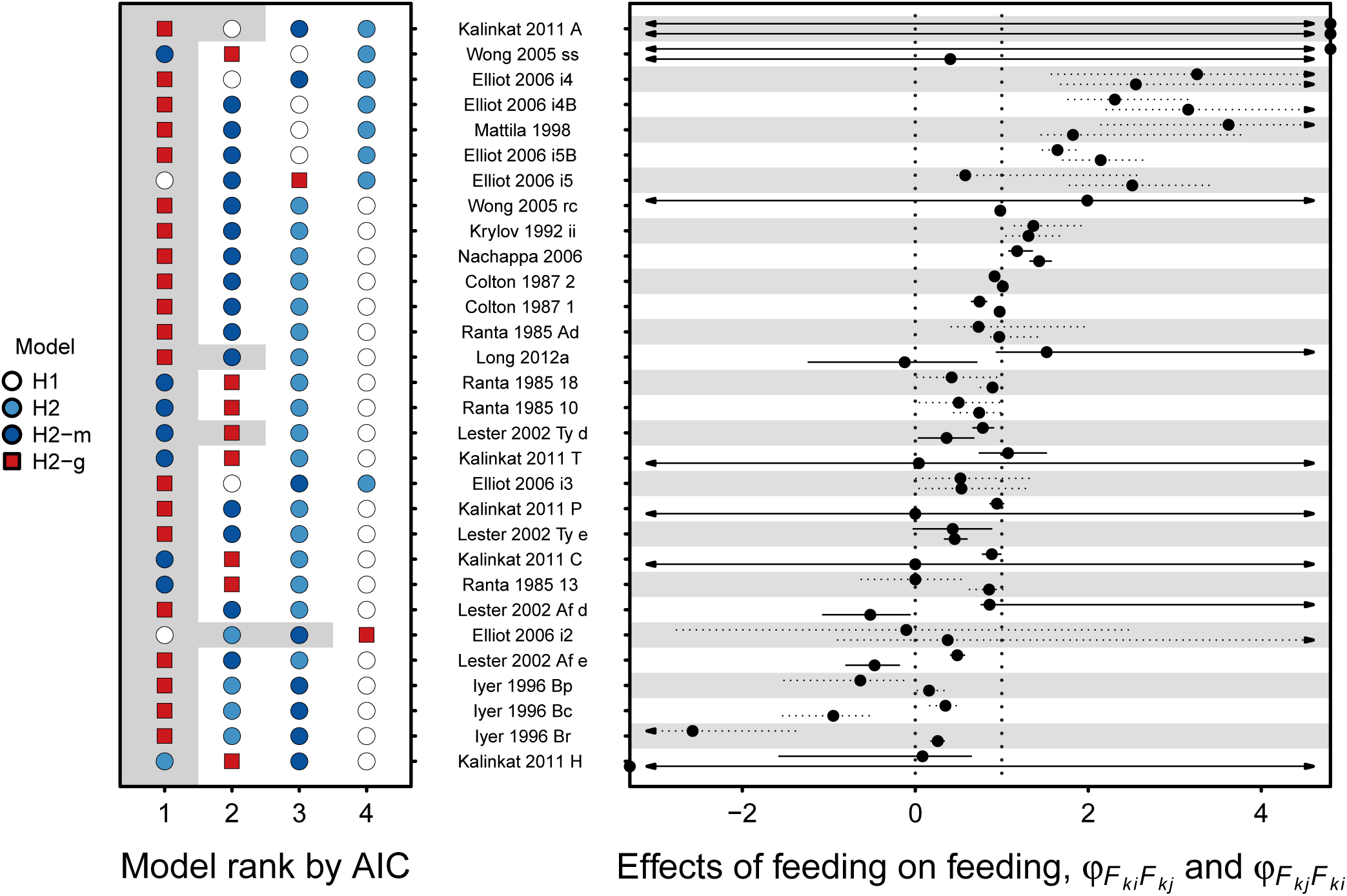
The effect that feeding on one prey species has on feeding on another as estimated across 30 multi-resource dependence datasets. One the left, we show the rank-order performance of functional-response models as judged by *AIC*, with rank 1 indicating the best model and rank 4 indicating the worst. The gray region indicates models with statistically equivalent support (i.e., Δ*AIC* < 2). The red square is the generalized multi-resource Holling Type II model introduced here (H2-g; Eq. 10). (See Table 2 for all model abbreviations.) On the right, we show the estimated mean and uncertainty for the effects feeding on one prey has on feeding on the other (i.e., 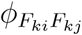 and 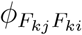 in Eq. 10). For visual clarity, the horizontal grey bands pair together the two parameter estimates corresponding to each individual dataset. Both points falling on the vertical line at 0 corresponds to the single-resource Holling Type II model (H2), and both points falling on the vertical line at 1 corresponds to the multi-resource Holling Type II model (H2-m). The large uncertainty for some points is most likely due to poor parameter identifiability. All line types are as in Fig. 2.

## Discussion

Our analyses provide compelling evidence that the processes affecting resource- and consumer dependence in feeding rates are frequently density dependent themselves. Across a large proportion of the single-resource consumer-dependence datasets, we observed that feeding and interference are rarely mutually exclusive. Likewise, we observed that a consumer’s behavior when feeding on one resource can appear very different to its behavior when feeding on another. That we were able to obtain these inferences despite the fact that, to our knowledge, the experimental design of none of the analyzed datasets was developed to measure our additional parameters, lends further credibility to our conclusions. We thus predict that evidence in support of functional-response models containing higher-order model terms will increase as datasets with larger sample sizes and targeted experimental designs are generated in the future.

Across the single-resource consumer-dependence datasets, the Beddington–DeAngelis model provided a reasonably good approximation to a rather large number of the single-resource consumer-interference datasets, even when that model was not statistically “best” (Fig. 2). In terms of their point estimates, the vast majority of these datasets had 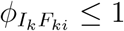, indicating that per capita feeding rates saturate at ever decreasing levels as the number of consumers increases. Phenomenologically, this arises either because the total time spent handling resources increases in higher consumer-density situations, or because consumers spend more and more time interfering with conspecifics that they would have otherwise spent feeding. In some datasets (e.g., “Chong 2006” and “Crowley 1989”), feeding in the presence of just over 3 additional conspecific consumers led to an effective doubling of each consumer’s handling time per resource consumed, relative to that of an isolated consumer individual. Dynamically, such “self-limitation” would lead to larger equilibrium resource densities and smaller equilibrium consumer densities. In contrast, a smaller subset of consumer-interference datasets suggest that consumers spent less and less time handling resources when feeding in the presence of more and more conspecifics (e.g., “Long 2012b” and “Kratina 2009”), which may be indicative of cooperative foraging.

Across the multi-resource dependence datasets, we found that most were inconsistent with the assumptions implied by either the single-resource or multi-resource Holling Type II functional responses, and hence neither classical model obtained widespread support. In both the empirical and theoretical literatures, it is common for researchers to decide a priori which model is most appropriate given their focal consumer and to analyze their data accordingly. Our results indicate that this may be an unwise path to follow since one would almost always need to know the characteristics of the consumers *and* resources before being able to adequately describe feeding rates. Moreover, a given consumer’s feeding rate could just as easily appear consistent with one model for a first resource and with a different model for another (e.g., “Long 2012a” where 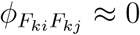 and 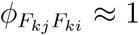), or somewhere in between (e.g., “Lester 2002 Ty d” where 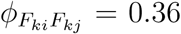 and 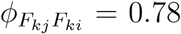). Clearly, more empirical research is needed to understand the biology that determines why each consumer–resource–resource combination lands in one particular location along the spectra of process inter-dependencies. The multi-resource Holling Type II is also widely used in simulations of food webs and other complex communities (Brose *et al*., 2006; Williams *et al*., 2007; Berlow *et al*., 2009; Iles & Novak, 2016; Delmas *et al*., 2017). Our results therefore challenge this assumption of theory as well, suggesting that ecological communities are most likely composed of a much broader array of consumer types. The exact dynamical properties that these varied consumer types may impart to their populations and food webs remain unknown. However, research on apparent competition (Holt, 1977; Abrams & Matsuda, 1996) and analyses of other functional-response models suggest their effects could be quite strong (Adamson & Morozov, 2013; 2014; Aldebert & Stouffer, 2018; Coblentz & DeLong, 2020).

### Functional responses emerge from independent and non-independent processes

Our analyses emphasize the fact that even the simplest functional responses are impacted by more than static attack rates, handling times, and interference rates. Instead, we argue it is more instructive to think about feeding as just one of multiple *processes* in which a predator could be engaging at any given moment of time (see also Koen-Alonso, 2007; Kéfi *et al*., 2012; Lafferty *et al*., 2015). The parameters 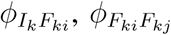, and/or 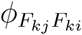 introduced here then allow us to quantify the first-order dependencies that exist between these processes. Importantly, while Crowley & Martin (1989) considered these dependencies to determine the extent to which a predator can or cannot interfere and feed simultaneously, we show here that viewing this as a dichotomy is vastly oversimplified. Instead, it is more appropriate to consider the various *ϕ*’s as capturing two key features of biological relevance: First, they allow us to statistically infer whether or not the rates of two processes proceed independently of each other. Second, when those rates do not proceed independently, they capture whether one process can be said to accelerate or decelerate the other. Within a functional-response context, we expect that this ontology will come rather naturally. After all, the Holling Type II functional response emerges precisely from the separation and assumed mutual exclusion of time spent searching and time spent handling (Holling, 1959b).

The lessons learned here likely apply to many other areas of ecology and biology as well. For example, the widespread use of Holling Type II functional responses and models like it in the study of plant–pollinator interactions (Holland *et al*., 2002; Morris *et al*., 2010) contrasts sharply with evidence that plant-neighborhood effects on pollinator behavior can be complex (Underwood *et al*., 2020). Models of such mutualisms often assume a priori that no interference occurs between pollinators (Okuyama & Holland, 2008; Vàzquez *et al*., 2015; Valdovinos, 2019). Based on our analyses, we expect there to be many more under-estimated processes at play in these systems, extending well beyond the densities of any focal interacting pair. Beyond consumer-resource interactions, standard growth-response curves (Tilman, 1977; 1982; Rothhaupt, 1988; Dybzinski & Tilman, 2007; Letten *et al*., 2018) and models of enzyme kinetics (Michaelis & Menten, 1913) applied to plant and microbial systems are both mathematical equivalents to the single-resource Holling Type II functional response, yet are routinely adopted in multi-resource contexts (Descamps-Julien & Gonzalez, 2005; Kleinhesselink & Adler, 2015; Letten *et al*., 2018). This occurs despite the tremendous utility that exists in identifying scenarios in which access to multiple resources synergistically promotes or retards growth (Sperfeld *et al*., 2012; Jeyasingh *et al*., 2020). Indeed, from trophic interactions and growth models to epistasis (Poelwijk *et al*., 2016; Sailer & Harms, 2017a;b) and drug-drug interactions (Tekin *et al*., 2018; Katzir *et al*., 2019), there are countless areas of biology in which researchers are interested in ways to quantify similar forms of non-independence and non-additivity. Our mathematical framework provides a general basis with which to explore each of these and others, following a tradition of embracing biological complexity rather than shying away from it (Evans *et al*., 2013).

As useful as this shift in perspective might appear, we nevertheless admit that it is not without practical limitations. As the numbers of processes and species under consideration increase, so too does the maximum number of potential parameters at a disproportionately high rate. In our “simple” scenario of a single consumer feeding on two resources, there were just two new parameters linking the two feeding rates; with three resources there are six new parameters, with four there are 12, and so on. The situation with multiple consumers *and* multiple resources becomes even more extreme. On the plus side, the data themselves can impose limits on model complexity since many processes of relevance to feeding rates—such as consumer interference and even prey handling—can only be measured indirectly in terms of their impact on feeding rates, and only one such implicit process can be measured per species density (Supplementary Material). Greater statistical power can therefore be achieved by obtaining information on additional response variables beyond feeding rates, as each such variable will contribute to the statistical likelihood of a given experimental replicate (Arditi & Glaizot, 1995). That said, the correct interpretation of such measured response variables is often not as cut and dry as it is with feeding (where resources are either consumed or not consumed), with even observable “handling times” not necessarily reflecting a rate-limiting process (Jeschke *et al*., 2002) and observable antagonistic encounters among individuals not necessarily reflecting rate-altering behavioral effects (Sheriff *et al*., 2020).

Given these practical limitations, a major challenge is to find model simplifications that can be supported, and to determine robust strategies for doing so. One option is to adopt a descriptive approach as proposed by Arditi & Michalski (1996); its phenomenological nature lends itself to a quick reduction in the number of parameters needing to be inferred. Alternatively, one can follow a statistical approach to the problem. For example, our fits to the multi-resource dependent datasets always treated both resources as functionally distinct (i.e., *a*_*ki*_ ≠ *a*_*kj*_ and *h*_*ki*_ ≠ *h*_*kj*_). Given a many-resource dataset, it may instead be advantageous to assess whether two or more resources are functionally equivalent in terms of model fit (Carrara *et al*., 2015; Ovaskainen *et al*., 2017). The statistical approach could also treat variation between resources as a statistical random effect (Ovaskainen & Soininen, 2011; Ovaskainen *et al*., 2016) or allow parameter variation to mirror phylogenetic or trait distances (Kalinkat *et al*., 2013). Clearly, proper inference of any such models will benefit from increased replication (Novak & Stouffer, 2020), as well as more robust and creative study designs (e.g., Dell *et al*., 2014; Novak *et al*., 2017; Uszko *et al*., 2017). Even so, not all phenomenological or statistical approaches are guaranteed to be logically consistent (Arditi & Michalski, 1996; Morozov & Petrovskii, 2013). All models should therefore be tested against criteria beyond fit and parsimony before they are applied, for example in population models (Malard *et al*., 2020; Moisset de Espanés *et al*., 2020).

## Conclusions

The study of biological models serves a variety of purposes (Evans *et al*., 2013; Otto & Rosales, 2020). We have focused here on the ability of generalizable models to fit observed variation in feeding rates across a large collection of empirical datasets. Rather than introduce additional, phenomenological parameters in the way that can occur with statistical methods like multiple regression (e.g., the inclusion of *m*-way interaction terms; Cox, 1984), our mathematical approach demonstrates why such interaction terms emerge: the inter-dependence of biological processes. Biological explanations for the large variation we observed across datasets remain to be determined. We therefore hope our study will also provide a fruitful starting point for a more mechanistic synthesis in the not too distant future.

## Acknowledgments

We thank Stella Uiterwaal and Gregor Kalinkat for providing their compiled bibliographies of published functional response studies. For generously providing their data, we thank Sven Bacher, Shane Blowes, Kevin Chan, Juang-Horng Chong, Will Cresswell, Malcolm Elliot, Gregor Fussmann, Fatemeh Ganjisaffar, Mark Hebblewhite, Tom Hossie, Gregor Kalinkat, Pavel Kratina, Birgit Lang, Phil Lester, Chris Long, Punya Nachappa, Anders Nilsson, Christina Prokopenko, Timothy Pusack, John Reeve, Thierry Spataro, Adrian Stier, John Vucetich, Will White, and Melisa Wong, as well as all authors who made their data publicly available in repositories. We further thank Peter Abrams, Roger Arditi, Thomas Hossie, Gregor Kalinkat, Pavel Kratina, Hao Ran Lai, Andrew Letten, Michelle Marraffini, and Rogini Runghen for suggestions that improved the manuscript. DBS acknowledges the support of the Marsden Fund Council, from New Zealand Government funding (grant 16-UOC-008) and a University of Canterbury Erskine grant.

## Author contributions

DBS and MN conceived of the project, the modeling framework, and contributed to the writing of analysis code; MN compiled the empirical data; DBS led the writing of the manuscript.

## Code and data availability

Code for all analyses, as well as most datasets, are available at https://github.com/stoufferlab/general-functional-responses. These and additional data sets have also been posted to online repositories per agreement with data contributors (see Tables S1 & S2), or were obtained from repositories to which they had previously been posted by the original authors.

