## Supporting Information for "Hidden layers of density dependence in consumer feeding rates"

### 1 Empirical datasets

Table S1: A summary of single-resource consumer dependence datasets. “Dataset” refers to the specific experiment from the study, and ‘-’ implies there was only one dataset available. “Nobs” indicates the sample size. “Replaced” refers to whether the consumed resource was replaced during the study, which dictated our use of a binomial versus a Poisson likelihood. “Consumer” refers to the whether the consumer was a predator or a parasitoid. “Raw” refers to whether we were able to use the raw data at the level of each treatment replicate, or whether we instead used means and associated uncertainty intervals to produce bootstrapped datasets. “Type” refers to whether the data was provided to us by the author, was obtained from an online repository, or was extracted from the publication. “Source” refers to the figures and tables from which the data where extracted.

| Study | Dataset | Nobs | Replaced | Consumer | Raw | Type | Source | Citation |
|---|---|---|---|---|---|---|---|---|
| Chan *et al.* (2017) | ch | 10 | Yes | Predator | Yes | Provided | - | Chan *et al.* (2017) |
| Chan *et al.* (2017) | cs | 10 | Yes | Predator | Yes | Provided | - | Chan *et al.* (2017) |
| Chan *et al.* (2017) | lh | 10 | Yes | Predator | Yes | Provided | - | Chan *et al.* (2017) |
| Chan *et al.* (2017) | ls | 10 | Yes | Predator | Yes | Provided | - | Chan *et al.* (2017) |
| Chant & Turnbull (1966) | - | 15 | No | Predator | Yes | Extracted | Table 2 | Novak & Stouffer (2020) |
| Chong & Oetting (2006) | - | 126 | Yes | Parasitoid | Yes | Provided | - | Chong (2020) |
| Crowley & Martin (1989) | - | 60 | Yes | Predator | No | Extracted | Fig. 2 | Novak & Stouffer (2020) |
| Edwards (1961) | nm | 97 | Yes | Parasitoid | Yes | Extracted | Tables 1, 2 & 3 | Novak & Stouffer (2020) |
| Edwards (1961) | ts1 | 75 | Yes | Parasitoid | Yes | Extracted | Tables 1, 2 & 3 | Novak & Stouffer (2020) |
| Edwards (1961) | ts2 | 17 | Yes | Parasitoid | Yes | Extracted | Tables 1, 2 & 3 | Novak & Stouffer (2020) |
| Elliott (2005) | i2 | 400 | Yes | Predator | Yes | Provided | - | Elliott (2020a) |
| Elliott (2005) | i3 | 400 | Yes | Predator | Yes | Provided | - | Elliott (2020a) |
| Elliott (2005) | i4 | 400 | Yes | Predator | Yes | Provided | - | Elliott (2020a) |
| Elliott (2005) | i5 | 400 | Yes | Predator | Yes | Provided | - | Elliott (2020a) |
| Eveleigh & Chant (1982) | aa | 111 | No | Predator | No | Extracted | Tables 4, 5 & 8 | Novak & Stouffer (2020) |
| Eveleigh & Chant (1982) | ad | 226 | No | Predator | No | Extracted | Tables 4, 5 & 8 | Novak & Stouffer (2020) |
| Eveleigh & Chant (1982) | ap | 267 | No | Predator | No | Extracted | Tables 4, 5 & 8 | Novak & Stouffer (2020) |
| Eveleigh & Chant (1982) | pa | 111 | No | Predator | No | Extracted | Tables 4, 5 & 8 | Novak & Stouffer (2020) |
| Eveleigh & Chant (1982) | pd | 278 | No | Predator | No | Extracted | Tables 4, 5 & 8 | Novak & Stouffer (2020) |
| Eveleigh & Chant (1982) | pp | 298 | No | Predator | No | Extracted | Tables 4, 5 & 8 | Novak & Stouffer (2020) |
| Fussmann *et al.* (2005) | - | 101 | Yes | Predator | Yes | Provided | - | Fussmann (2020) |
| Griffen & Delaney (2007) | f1a | 108 | No | Predator | No | Extracted | Fig 1 & Fig A1 | Novak & Stouffer (2020) |
| Griffen & Delaney (2007) | f1b | 108 | No | Predator | No | Extracted | Fig 1 & Fig A1 | Novak & Stouffer (2020) |
| Griffen & Delaney (2007) | fA1a | 32 | No | Predator | No | Extracted | Fig 1 & Fig A1 | Novak & Stouffer (2020) |
| Griffen & Delaney (2007) | fA1b | 32 | No | Predator | No | Extracted | Fig 1 & Fig A1 | Novak & Stouffer (2020) |
| Hassan (1976) | Ag | 100 | Yes | Parasitoid | No | Extracted | Table 1 | Novak & Stouffer (2020) |
| Hassan (1976) | Br | 100 | Yes | Parasitoid | No | Extracted | Table 1 | Novak & Stouffer (2020) |
| Hassan (1976) | Pp | 100 | Yes | Parasitoid | No | Extracted | Table 1 | Novak & Stouffer (2020) |
| Hossie & Murray (2016) | cl | 42 | No | Predator | Yes | Provided | - | Hossie & Murray (2020) |
| Hossie & Murray (2016) | ev | 42 | No | Predator | Yes | Provided | - | Hossie & Murray (2020) |
| Huffaker & Matsumoto (1982) | - | 40 | Yes | Parasitoid | No | Extracted | Table 1 | Novak & Stouffer (2020) |
| Johnson (2006) | - | 45 | No | Predator | Yes | Extracted | Fig. 1 | Novak & Stouffer (2020) |
| Jones & Hassell (1988); Jones (1986) | e4 | 125 | No | Parasitoid | Yes | Extracted | Fig. 1a & 3 | Novak & Stouffer (2020) |
| Jones & Hassell (1988); Jones (1986) | e5 | 140 | No | Parasitoid | Yes | Extracted | Fig. 1a & 3 | Novak & Stouffer (2020) |
| Katz (1985) | - | 112 | No | Predator | No | Extracted | Table 1 | Arditi & Akçakaya (1990) |
| Kfir (1983) | - | 120 | No | Parasitoid | No | Extracted | Table 1 | Novak & Stouffer (2020) |
| Kratina *et al.* (2009) | - | 67 | No | Predator | Yes | Provided | - | Kratina (2020) |

Table S1: *(continued)*

| Study | Dataset | Nobs | Replaced | Consumer | Raw | Type | Source | Citation |
|---|---|---|---|---|---|---|---|---|
| Krylov (1992) | i | 28 | No | Predator | No | Extracted | Table 1 & Fig. 1L | Novak & Stouffer (2020) |
| Kumar & Tripathi (1985) | Cc | 90 | No | Parasitoid | No | Extracted | Tables 1 & 2 & Figs. 1 & 2 | Novak & Stouffer (2020) |
| Kumar & Tripathi (1985) | Dl | 90 | No | Parasitoid | No | Extracted | Tables 1 & 2 & Figs. 1 & 2 | Novak & Stouffer (2020) |
| Kumar & Tripathi (1985) | Sm | 90 | No | Parasitoid | No | Extracted | Tables 1 & 2 & Figs. 1 & 2 | Novak & Stouffer (2020) |
| Lang *et al.* (2012) | Po 10 | 185 | No | Predator | Yes | Provided | - | Lang (2020) |
| Lang *et al.* (2012) | Po 20 | 181 | No | Predator | Yes | Provided | - | Lang (2020) |
| Lang *et al.* (2012) | Pt 10 | 184 | No | Predator | Yes | Provided | - | Lang (2020) |
| Lang *et al.* (2012) | Pt 20 | 186 | No | Predator | Yes | Provided | - | Lang (2020) |
| Long *et al.* (2012b) | - | 50 | No | Predator | Yes | Provided | - | Long (2020a) |
| Mansour & Lipcius (1991) | - | 36 | No | Predator | No | Extracted | Fig. 1a | Novak & Stouffer (2020) |
| Médoc *et al.* (2013) | - | 48 | Yes | Predator | Yes | Provided | - | Médoc *et al.* (2020b) |
| Médoc *et al.* (2015) | be | 76 | Yes | Predator | Yes | Provided | - | Médoc *et al.* (2020a) |
| Médoc *et al.* (2015) | dv | 76 | Yes | Predator | Yes | Provided | - | Médoc *et al.* (2020a) |
| Médoc *et al.* (2015) | pu | 76 | Yes | Predator | Yes | Provided | - | Médoc *et al.* (2020a) |
| Mertz & Davies (1968) | - | 120 | No | Predator | Yes | Extracted | Table 1 | Novak & Stouffer (2020) |
| Mills & Lacan (2004) | - | 179 | Yes | Parasitoid | Yes | Extracted | Fig. 1a-c | Novak & Stouffer (2020) |
| Montoya *et al.* (2000) | - | 528 | Yes | Parasitoid | No | Extracted | Table 1 | Novak & Stouffer (2020) |
| Omkar & Pervez (2004) | - | 90 | No | Predator | Yes | Provided | - | Omkar & Pervez (2004) |
| Prokopenko *et al.* (2017) | - | 80 | Yes | Predator | Yes | Provided | - | Prokopenko (2020) |
| Pusack *et al.* (2018) | - | 60 | Yes | Predator | Yes | Provided | - | Pusack (2020) |
| Reeve (1997) | - | 26 | No | Predator | Yes | Provided | - | Reeve (2020) |
| Salt (1974) | - | 50 | Yes | Predator | No | Extracted | Fig. 3 & Table 1 | Novak & Stouffer (2020) |
| Uttley (1980) | i1 | 168 | No | Predator | No | Extracted | Figs. 4.3.1, 4.3.2, 4.3.3, 8.3.1 & 8.3.2 | Novak & Stouffer (2020) |
| Uttley (1980) | i2 | 204 | No | Predator | No | Extracted | Figs. 4.3.1, 4.3.2, 4.3.3, 8.3.1 & 8.3.2 | Novak & Stouffer (2020) |
| Uttley (1980) | i3 | 224 | No | Predator | No | Extracted | Figs. 4.3.1, 4.3.2, 4.3.3, 8.3.1 & 8.3.2 | Novak & Stouffer (2020) |
| Uttley (1980) | n1 | 266 | No | Predator | No | Extracted | Figs. 4.3.1, 4.3.2, 4.3.3, 8.3.1 & 8.3.2 | Novak & Stouffer (2020) |
| Uttley (1980) | n2 | 312 | No | Predator | No | Extracted | Figs. 4.3.1, 4.3.2, 4.3.3, 8.3.1 & 8.3.2 | Novak & Stouffer (2020) |
| Vahl *et al.* (2005) | k | 10 | No | Predator | Yes | Extracted | Fig. 4 | Novak & Stouffer (2020) |
| Vahl *et al.* (2005) | t | 10 | No | Predator | Yes | Extracted | Fig. 4 | Novak & Stouffer (2020) |
| Von Westernhagen & Rosenthal (1976) | 2hr | 80 | No | Predator | No | Extracted | Fig. 3 | Novak & Stouffer (2020) |
| Von Westernhagen & Rosenthal (1976) | 4hr | 120 | No | Predator | No | Extracted | Fig. 3 | Novak & Stouffer (2020) |
| Von Westernhagen & Rosenthal (1976) | 8hr | 100 | No | Predator | No | Extracted | Fig. 3 | Novak & Stouffer (2020) |
| Vucetich *et al.* (2002) | m14 | 118 | Yes | Predator | Yes | Provided | - | Vucetich *et al.* (2002); Jost *et al.* (2005) |
| Vucetich *et al.* (2002) | m98 | 77 | Yes | Predator | Yes | Provided | - | Vucetich *et al.* (2002); Jost *et al.* (2005) |
| Vucetich *et al.* (2002) | w14 | 44 | Yes | Predator | Yes | Provided | - | Vucetich *et al.* (2002); Jost *et al.* (2005) |
| Vucetich *et al.* (2002) | w98 | 28 | Yes | Predator | Yes | Provided | - | Vucetich *et al.* (2002); Jost *et al.* (2005) |
| Walde & Davies (1984) | - | 60 | Yes | Predator | No | Extracted | Fig 2 & 4 | Novak & Stouffer (2020) |
| Wasserman *et al.* (2016b) | bg | 38 | No | Predator | Yes | Repository | - | Wasserman *et al.* (2016a) |
| Wasserman *et al.* (2016b) | mb | 37 | No | Predator | Yes | Repository | - | Wasserman *et al.* (2016a) |
| Wasserman *et al.* (2016b) | ti | 39 | No | Predator | Yes | Repository | - | Wasserman *et al.* (2016a) |

Table S2: A summary of multi-resource dependence datasets. “Dataset” refers to the specific experiment from the study, and ‘-’ implies there was only one dataset available. “Nobs” indicates the sample size per resource consumed. “Replacement” refers to whether the consumed resources were replaced during the study, which dictated our use of a binomial versus a Poisson likelihood. “Consumer” refers to whether the consumer was a predator or a parasitoid. “Raw” refers to whether we were able to use the raw data at the level of each treatment replicate, or whether we instead used means and associated uncertainty intervals to produce bootstrapped datasets. “Type” refers to whether the data was provided to us by the author, was obtained from an online repository, or was extracted from the publication. “Source” refers to the figures and tables from which the data where extracted.

| Study | Dataset | Nobs | Replaced | Consumer | Raw | Type | Source | Citation |
|---|---|---|---|---|---|---|---|---|
| Colton (1983; 1987) | 1 | 108 | No | Predator | Yes | Extracted | Table B3 | Novak & Stouffer (2020) |
| Colton (1983; 1987) | 2 | 108 | No | Predator | Yes | Extracted | Table B3 | Novak & Stouffer (2020) |
| Elliott (2006) | i2 | 290 | Yes | Predator | No | Provided | - | Elliott (2020b) |
| Elliott (2006) | i3 | 290 | Yes | Predator | No | Provided | - | Elliott (2020b) |
| Elliott (2006) | i4 | 290 | Yes | Predator | No | Provided | - | Elliott (2020b) |
| Elliott (2006) | i4B | 290 | Yes | Predator | No | Provided | - | Elliott (2020b) |
| Elliott (2006) | i5 | 290 | Yes | Predator | No | Provided | - | Elliott (2020b) |
| Elliott (2006) | i5B | 290 | Yes | Predator | No | Provided | - | Elliott (2020b) |
| Iyer & Rao (1996) | Bc | 192 | No | Predator | No | Extracted | Fig. 1 & 2 | Novak & Stouffer (2020) |
| Iyer & Rao (1996) | Bp | 192 | No | Predator | No | Extracted | Fig. 1 & 2 | Novak & Stouffer (2020) |
| Iyer & Rao (1996) | Br | 192 | No | Predator | No | Extracted | Fig. 1 & 2 | Novak & Stouffer (2020) |
| Kalinkat *et al.* (2011) | A | 48 | Yes | Predator | Yes | Provided | - | Kalinkat *et al.* (2018) |
| Kalinkat *et al.* (2011) | C | 49 | Yes | Predator | Yes | Provided | - | Kalinkat *et al.* (2018) |
| Kalinkat *et al.* (2011) | H | 48 | Yes | Predator | Yes | Provided | - | Kalinkat *et al.* (2018) |
| Kalinkat *et al.* (2011) | P | 70 | Yes | Predator | Yes | Provided | - | Kalinkat *et al.* (2018) |
| Kalinkat *et al.* (2011) | T | 137 | Yes | Predator | Yes | Provided | - | Kalinkat *et al.* (2018) |
| Krylov (1992) | ii | 56 | No | Predator | No | Extracted | Fig. 1 & Fig. 2 | Novak & Stouffer (2020) |
| Lester & Harmsen (2002) | Af d | 60 | Yes | Predator | Yes | Provided | - | Lester (2020) |
| Lester & Harmsen (2002) | Af e | 60 | Yes | Predator | Yes | Provided | - | Lester (2020) |
| Lester & Harmsen (2002) | Ty d | 60 | Yes | Predator | Yes | Provided | - | Lester (2020) |
| Lester & Harmsen (2002) | Ty e | 60 | Yes | Predator | Yes | Provided | - | Lester (2020) |
| Long *et al.* (2012a) | - | 94 | No | Predator | Yes | Provided | - | Long (2020b) |
| Mattila & Bonsdorff (1998) | - | 37 | No | Predator | No | Extracted | Fig. 3a & 4 | Novak & Stouffer (2020) |
| Nachappa *et al.* (2006) | - | 158 | No | Predator | Yes | Provided | - | Nachappa *et al.* (2006) |
| Ranta & Nuutinen (1985) | 10 | 123 | Yes | Predator | No | Extracted | Fig. 2 & 3 & Tables 2 & 5 | Novak & Stouffer (2020) |
| Ranta & Nuutinen (1985) | 13 | 123 | Yes | Predator | No | Extracted | Fig. 2 & 3 & Tables 2 & 5 | Novak & Stouffer (2020) |
| Ranta & Nuutinen (1985) | 18 | 123 | Yes | Predator | No | Extracted | Fig. 2 & 3 & Tables 2 & 5 | Novak & Stouffer (2020) |
| Ranta & Nuutinen (1985) | Ad | 123 | Yes | Predator | No | Extracted | Fig. 2 & 3 & Tables 2 & 5 | Novak & Stouffer (2020) |
| Wong & Barbeau (2005) | rc | 48 | Yes | Predator | Yes | Provided | - | Wong & Barbeau (2020) |
| Wong & Barbeau (2005) | ss | 48 | Yes | Predator | Yes | Provided | - | Wong & Barbeau (2020) |

### 2 Maximum-likelihood parameter estimation

For both single-resource and multiple-resource datasets in which consumed resources were continually replaced, we assumed that observed *counts* of resources consumed were drawn from a Poisson distribution parameterized by the feeding rate of the functional response. For datasets in which consumed resources were not replaced, we assumed that observed *proportions* of resources consumed were drawn from a binomial distribution parameterized by the feeding rate of the functional response and by the total number of available resources at the start of the experiment. For non-replacement datasets with a single resource, we generated analytical predictions for the proportion of resources consumed using the Lambert W function to account for continuous resource depletion (Okuyama & Ruyle, 2011; Lehtonen, 2016; Rosenbaum & Rall, 2018). For non-replacement datasets with multiple resources, we determined the proportion of both resources consumed via numerical integration of the coupled differential equations (Keitt, 2017). For multiple-resource datasets the likelihood was calculated as the sum of the likelihoods of each of the two resources. This is because the functional response parameters for each resource influence both feeding rates (except in the limiting cases) and hence require simultaneous estimation.

### 3 Parametric bootstrapping of empirical datasets

For each dataset comprised of treatment means and associated uncertainties, we generated parametrically-bootstrapped datasets as follows. For a treatment level consisting of $n$ replicates with mean number of resources consumed $\mu$ and standard error $s$, we first drew $n$ random values $x$ from a Gaussian distribution with mean $\mu$ and standard deviation $\sigma = s\sqrt{n}$. To mirror our inference process and ensure data consistent with the underlying likelihood functions, we then used these random values to parameterize Poisson or binomial processes depending on whether the original dataset corresponded to a replacement or non-replacement study design, respectively. For Katz (1985) where we could not obtain estimates of uncertainty, we used the available means as raw data.

To maintain biologically sensible predictions, we replaced any random value $x < 0$ with 0 for both experiment types since this is the lower limit of observable feeding. In non-replacement datasets, we replaced any value of $x > N$ with $N$ since this is the upper limit of observable feeding. For the multi-resource datasets, we were able to obtain uncertainties of feeding on both resources but not the covariance between them. We therefore simulated these paired observations as independent random variables.

### 4 Model comparison based on $AIC_C$ and $BIC$

*AIC* is the most frequently used information criterion for evaluating relative model performance (Aho *et al.*, 2014). It estimates the expected relative Kullback–Leibler divergence of a focal model from the true model responsible for generating the data. A second information criterion, $AIC_C$, incorporates a correction for the bias that *AIC* exhibits at small sample sizes, imposing a greater penalty on a model's parametric complexity than does *AIC*. The difference between *AIC* and $AIC_C$ diminishes as sample size increases. However, the standard equation for $AIC_C$ was derived in a context of linear regression models with Gaussian errors and hence is a biased estimator for non-linear models of the sort analyzed here (Burnham & Anderson, 2002). A third criterion is the Bayesian Information Criterion, *BIC* (a.k.a. Schwarz Information Criterion; Schwarz, 1978). Unlike for *AIC* and $AIC_C$, a model's parametric complexity as estimated by *BIC* increases with sample size $n$. Among their different underlying assumptions, *BIC* presumes the true model to be among the considered models while *AIC* does not. Pragmatically speaking, *BIC* penalizes complex models to a greater extent than does *AIC* (when sample size is greater than 8) and therefore favors models with fewer parameters. We examined whether the model comparisons presented in the main text depended on our choice of *AIC* as our baseline criterion and found that the results were qualitatively similar regardless of which information criterion was applied (Figs. S1 & S2).

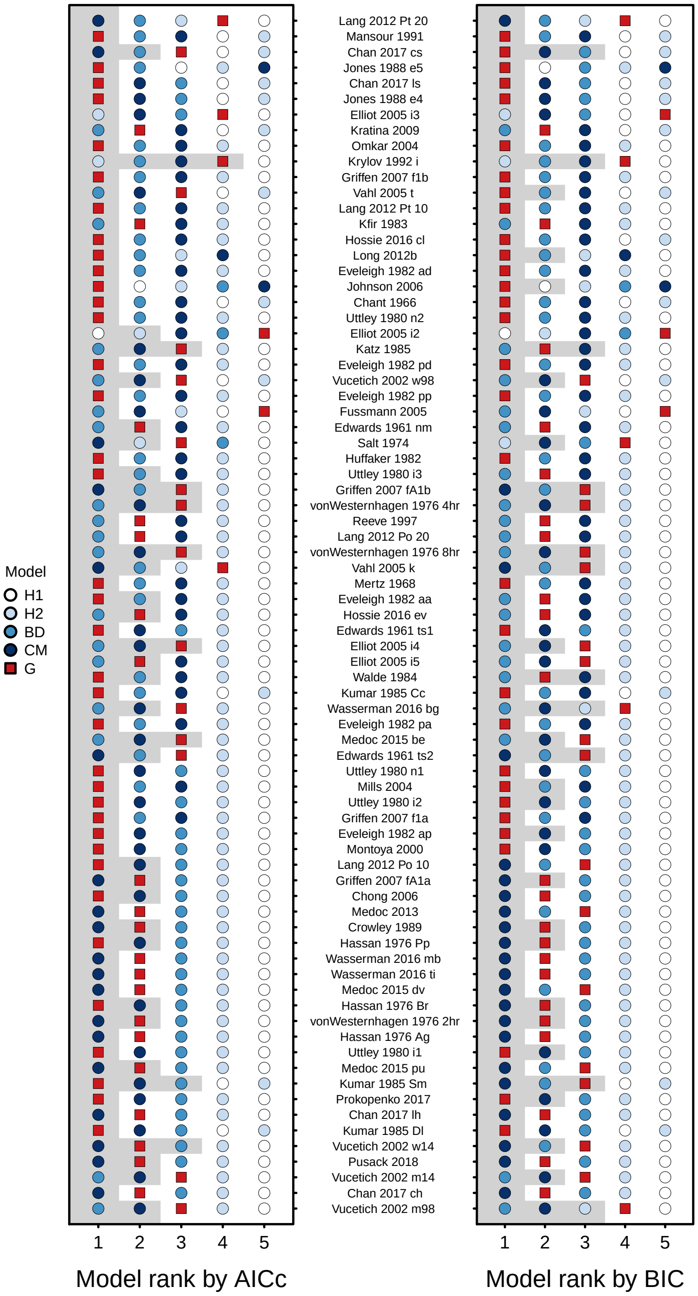


Figure S1: Model comparison across 77 single-resource consumer-dependence datasets with alternative information criteria. One the left, we show the rank-order performance of functional-response models as judged by $AIC_C$, with rank 1 indicating the best model and rank 5 indicating the worst. One the right, we show the rank-order performance of functional-response models as judged by $BIC$, with rank 1 again indicating the best model and rank 5 indicating the worst. The gray region in both plots indicates models with statistically equivalent support (i.e., $\Delta IC < 2$). The red square is the generalized consumer-dependent model introduced in the main text (G).

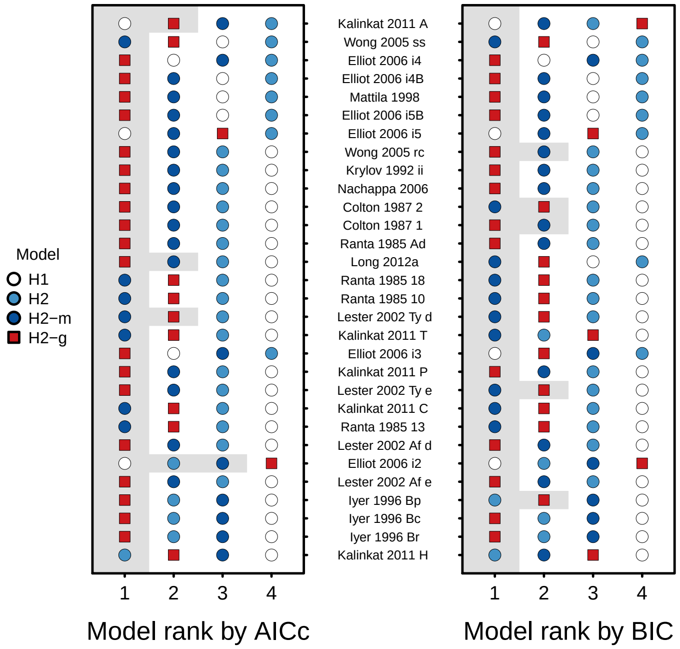


Figure S2: Model comparison across 30 multi-resource dependence datasets with alternative information criteria. One the left, we show the rank-order performance of functional-response models as judged by $AIC_C$, with rank 1 indicating the best model and rank 5 indicating the worst. One the right, we show the rank-order performance of functional-response models as judged by $BIC$, with rank 1 again indicating the best model and rank 5 indicating the worst. The gray region in both plots indicates models with statistically equivalent support (i.e., $\Delta IC < 2$). The red square is the generalized consumer-dependent model introduced in the main text (G).

### 5 Limit of one implicit density-dependent process per predictor density

Consider a single consumer $k$ foraging on individuals from a single resource species $i$, and experimental data which provide estimates of feeding rate $F_{ki}$ as a function of varying density of resource $N_i$. In this case, our functional response can only be a linear or non-linear function of a single predictor density: $N_i$. Rather than a single "handling time" quantified with the parameter $h_{ki}$ as occurs in a Holling Type II model, imagine that we also wish to separate handling into "capture time" $\gamma_{ki}$ and "digestion time" $\delta_{ki}$ (Jeschke *et al.*, 2002). Similar to the main text, we can define the feeding rates $F_{ki}$ as

$$F_{ki}\left(N_i\right) \quad = \quad a_{ki}N_i\left(1-\gamma_{ki}F_{ki}-\delta_{ki}F_{ki}\right)\ . \tag{S1}$$

If we rearrange this equation algebraically to solve for $F_{ki}$, we obtain

$$F_{ki}\left(N_i\right) \quad = \quad \frac{a_{ki}N_i}{1+a_{ki}\left(\gamma_{ki}+\delta_{ki}\right)N_i}\ , \tag{S2}$$

which implies that resource-dependent variation in feeding rates alone is insufficient for us to distinguish between capture and digestion in the absence of additional, independent information with which to estimate one or both of the parameters $\gamma_{ki}$ and $\delta_{ki}$.

Now consider a dataset with varying densities $N_k$ of the consumer $k$ foraging on varied densities $N_i$ of a single resource species $i$. In this case, our functional response can only be a linear or non-linear function of two predictor densities: $N_i$ and $P_k$. Rather than a single interference process occurring between conspecific consumers, we wish to distinguish between direct antagonistic interactions between consumers, which has an associated "antagonism" time $\xi_k$ (during which consumers cannot feed), and avoidance of interactions between consumers, which has an associated "avoidance" time $\kappa_k$ (also during which consumers cannot feed). Similar to the main text, we can define the feeding rate $F_{ki}$, the interference rate $I_k$, and the avoidance rate $A_k$ as

$$\begin{aligned}
I_k\left(N_i,P_k\right) &= \alpha_k(P_k-1)\left(1-h_{ki}F_{ki}-\kappa_kA_k-\xi_kI_k\right) && \text{(S3)}\\
A_k\left(N_i,P_k\right) &= \eta_k(P_k-1)\left(1-h_{ki}F_{ki}-\kappa_kA_k-\xi_kI_k\right) && \text{(S4)}\\
F_{ki}\left(N_i,P_k\right) &= a_{ki}N_i\left(1-h_{ki}F_{ki}-\kappa_kA_k-\xi_kI_k\right)\ . && \text{(S5)}
\end{aligned}$$

If we rearrange this equation algebraically to solve for $F_{ki}$, we obtain

$$F_{ki}\left(N_i, P_k\right) = \frac{a_{ki} N_i}{1 + a_{ki} h_{ki} N_i + \left(\alpha_k \xi_k + \eta_k \kappa_k\right)\left(P_k - 1\right)}\,, \tag{S6}$$

which implies that resource- and consumer-dependent variation in feeding rates is insufficient for us to distinguish between antagonism and avoidance in the absence of additional, independent information about one or both of $I_k$ and $A_k$.

Another way to understand this limitation is to consider the statistical identifiability of the parameters in the functional responses given by Eq. (S2) & (S6). In order for a set of parameters to be identifiable, the derivatives of the generating function with respect to those parameters should all be unique (Beck & Arnold, 1977). This does not hold true for either of the examples above; for example, looking at Eq. (S2)

$$\frac{\partial F_{ki}}{\partial \gamma_{ki}} = \frac{\partial F_{ki}}{\partial \delta_{ki}} = \frac{-\left(a_{ki} N_i\right)^2}{\left(1 + a_{ki}\left(\gamma_{ki} + \delta_{ki}\right) N_i\right)^2}\,. \tag{S7}$$

A somewhat counter-intuitive consequence of this is that the limit of one implicit process per predictor density holds only so long as the order of that density remains the same, where order reflects the implicit or explicit exponent of the predictor density (e.g., $N_i$ is order $1$, $(N_i)^2$ is order $2$, and so on). Therefore, one cannot separate capture time from digestion time in our example because both are order $(N_i)^1$ processes in that they occur *per resource consumed*. If one process was order $1$ (i.e., depended on $(N_i)^1$) and the other was order $2$ (i.e., depended on $(N_i)^2$), both *could* be inferred simultaneously. Indeed, this is the reason why we can infer additional parameters that are multiplicative *combinations* of predictor densities in the models introduced here and also the reason why it is possible to shift from a linear Holling Type I model to the non-linear Holling Type II model when we have a single order $1$ predictor density $N_i$.

### 6 Multi-resource dependence and prey electivities

In a multi-resource context, it is common to think about variation in feeding rates across resources as being driven by "prey electivities" or "predator preferences" (Ivlev, 1961; Chesson, 1983; 1989; Murdoch, 1969; Oaten & Murdoch, 1975). Given a choice of multiple resources (prey), the optimal foraging strategy of a given consumer (predator) may vary depending on resource availability. For example, a high feeding rate on of one type of resource due to that resource being abundant may lead to the consumer feeding differentially on another resource to achieve nutritional balance. In a functional-response context, one way to determine preferences is based on the ratio of attack rates. From this perspective, our generalized multi-resource Type II functional response (Eq. 10) is consistent with consumers having fixed preferences because the attack rates are constant. Analytically, this constancy of attack rates can also be seen by virtue of the fact that there is no common factor with which any $a_{ki}$ is multiplied in both the numerator *and* denominator of Eq. (10).

That said, the ratio of attack rates only fully captures realized preferences and realized differences in relative consumption when handling times are negligible. One can alternatively quantify preferences and/or switching behavior based on the proportion of resources consumed relative to the proportion of resources available (i.e., $F_{ki}/F_{kj}$ versus $N_i/N_j$ (Murdoch, 1969; van Leeuwen *et al.*, 2013). For Eq. (10) from the main text, we thereby obtain

$$\frac{F_{ki}}{F_{kj}} = \frac{a_{ki}\left[1+\left(1-\phi_{F_{ki}F_{kj}}\right)a_{kj}h_{kj}N_j\right]}{a_{kj}\left[1+\left(1-\phi_{F_{kj}F_{ki}}\right)a_{ki}h_{ki}N_i\right]} \times \frac{N_i}{N_j}\,. \tag{S8}$$

This perspective allows us to see how non-independent feeding (and the parameters $\phi_{F_{ki}F_{kj}}$ and $\phi_{F_{ki}F_{kj}}$) induce apparent switching behavior because the proportionality constant in front of $\frac{N_i}{N_j}$ is only constant when $\phi_{F_{ki}F_{kj}} = \phi_{F_{ki}F_{kj}} = 1$. An equivalent inference is obtained when quantifying preferences by normalizing resource-specific foraging by the total foraging (i.e., $F_{ki}/(F_{ki} + F_{kj})$). With the caveat that the multiple-resource datasets in our collection are primarily laboratory-based, this scenario appears to be rare (see Fig. 3 and the estimates of $\phi_{F_{ki}F_{kj}}$ and $\phi_{F_{ki}F_{kj}}$).

Conspicuously absent from Eq. (S8) are $N_i$ in the numerator and $N_j$ in the denominator of the proportionality constant (cf. van Leeuwen *et al.*, 2013). Such density dependence can be obtained by modifying Eqs. (8) & (9) of the main text to make the attack rates explicit functions of the other resources' density. Although this phenomenological manipulation of the model stands in contrast to the way density-dependent handling times "emerge" as shown in the main text, it nonetheless provides a useful basis of comparison.

For example, we could include linearly-dependent attack rates as

$$\tilde{F}_{ki}\left(N_i, N_j\right) = \tilde{a}_{ki} N_i \left(1 - h_{ki}\tilde{F}_{ki} - \phi_{\tilde{F}_{ki}\tilde{F}_{kj}} h_{kj}\tilde{F}_{kj}\right) \quad \text{(S9)}$$

$$\tilde{F}_{kj}\left(N_i, N_j\right) = \tilde{a}_{kj} N_j \left(1 - \phi_{\tilde{F}_{kj}\tilde{F}_{ki}} h_{ki}\tilde{F}_{ki} - h_{kj}\tilde{F}_{kj}\right) \quad \text{(S10)}$$

$$\tilde{a}_{ki} = a_{ki}\left(1 + \alpha_{kij} N_j\right) \quad \text{(S11)}$$

$$\tilde{a}_{kj} = a_{kj}\left(1 + \alpha_{kji} N_i\right) , \quad \text{(S12)}$$

introducing two new parameters, $\alpha_{kij}$ and $\alpha_{kji}$, that control the effect of the abundance of one resource on the consumer's attack rate on the other resource. Since these changes directly modify the attack rates, they can be carried through upon solving for $\tilde{F}_{ki}$ and $\tilde{F}_{kj}$, just as shown in the main text. The resulting expression for $\tilde{F}_{ki}$ is

$$\tilde{F}_{ki}\left(N_i, N_j\right) = \frac{\tilde{a}_{ki} N_i \left[1 + \left(1 - \phi_{\tilde{F}_{ki}\tilde{F}_{kj}}\right) \tilde{a}_{kj} h_{kj} N_j\right]}{\left(1 + \tilde{a}_{ki} h_{ki} N_i\right)\left(1 + \tilde{a}_{kj} h_{kj} N_j\right) - \phi_{\tilde{F}_{ki}\tilde{F}_{kj}} \phi_{\tilde{F}_{kj}\tilde{F}_{ki}} \tilde{a}_{ki} h_{ki} \tilde{a}_{kj} h_{kj} N_i N_j} , \quad \text{(S13)}$$

which is equivalent to Eq. (10) but with $a_{ki}$ replaced by $\tilde{a}_{ki}$ and $a_{kj}$ replaced by $\tilde{a}_{kj}$. The corresponding expression for predator preferences (defined as above) is

$$\frac{\tilde{F}_{ki}}{\tilde{F}_{kj}} = \frac{a_{ki}\left(1 + \alpha_{kij} N_j\right)\left[1 + \left(1 - \phi_{\tilde{F}_{ki}\tilde{F}_{kj}}\right) a_{kj}\left(1 + \alpha_{kji} N_i\right) h_{kj} N_j\right]}{a_{kj}\left(1 + \alpha_{kji} N_i\right)\left[1 + \left(1 - \phi_{\tilde{F}_{kj}\tilde{F}_{ki}}\right) a_{ki}\left(1 + \alpha_{kij} N_j\right) h_{ki} N_i\right]} \times \frac{N_i}{N_j} . \quad \text{(S14)}$$

Notably, the proportionality constant in this expression includes $N_i$ in the numerator and $N_j$ in the denominator. These densities are *always* multiplied by the density of the "other" resource. This implies that the only way to have these densities appear alone is to introduce them in our expressions for density-dependent attack rates (Eqs. S11 & S12). (Note that the functional form of density-dependent preferences achieved phenomenologically in Eq. (S13) and captured by Eq. (S8) are *identical* when $\phi_{F_{ki}F_{kj}} = \phi_{F_{ki}F_{kj}} = 1$.)

As might be expected, including density-dependent attack rates *and* non-independent feeding (via $\phi_{F_{ki}F_{kj}}$ and $\phi_{F_{kj}F_{ki}}$) creates many additional higher-order terms in the resulting functional responses. That is, the numerator and denominator of Eq. (S13) could include terms in the numerator and denominator of order 2 that depend on $(N_i)^2$, $(N_j)^2$, $(N_i)^2(N_j)^1$, $(N_i)^1(N_j)^2$, etc. The only exception is when feeding on each resource is 100% independent of feeding on the other (i.e., $\phi_{F_{ki}F_{kj}} = \phi_{F_{ki}F_{kj}} = 0$), in which case the orders of Eq. (S13) is equivalent to those of Eq. (10). Again with the caveat that the multiple-resource datasets in our empirical collection are primarily laboratory-based, this scenario also appears to be rare (see Fig. 3 and the estimates of $\phi_{F_{ki}F_{kj}}$ and $\phi_{F_{ki}F_{kj}}$).
